# Expanding Hippocampal Ensembles through LINE-1 RNA Downregulation Enhances Cognition

**DOI:** 10.64898/2026.10.14.758979

**Authors:** Nicole Politowska, Judit Gal Toth, Peter U. Hámor, Irena Pigulevskiy, Jacqueline P. Welday, Faten Taki, Hiba Nawaid, Rose Sciortino, Kristen Pleil, Miklos Toth

## Abstract

Episodic memories are encoded through the activation of small overlapping neuronal ensembles, a coding principle that ensures efficient encoding and high memory capacity. While cellular excitability is known to drive the allocation of select hippocampal neurons into these ensembles, the underlying molecular mechanisms remain elusive. Here, an epigenetic-functional analysis of hippocampal dentate granule cells revealed robust nuclear expression of RNA from autonomous LINE-1 retrotransposons, with significantly lower LINE-1 RNA levels in environmentally activated neurons compared to non-activated counterparts. Knockdown of LINE-1 RNA in the dentate gyrus increased action potential firing capacity and expanded the number of environmentally activated cells, indicating that LINE-1 expression is inversely correlated with ensemble eligibility. LINE-1 RNA knockdown enhanced spatial working memory at the expense of pattern separation, a trade-off consistent with expanded dentate gyrus ensembles and diminished sparse coding. Together, these findings demonstrate that nuclear LINE-1 RNA was co-opted during evolution to control ensemble eligibility in dentate granule cells, a novel postmitotic neuronal function for LINE-1 RNA that restricts ensemble size to preserve sparse coding and support pattern separation.

---

Encoding of hippocampal episodic memories relies on the activation of small ensembles of overlapping neurons, a high-capacity mechanism that ensures an efficient and specific representation of individual experiences. Restriction in neuronal recruitment is most pronounced within the dentate gyrus (DG), wherein only a few percent of dentate granule cells (DGCs) are allocated to an ensemble^1,2^. This sparse recruitment allows the DG to execute pattern separation—the orthogonalization of highly similar environmental contexts—while concurrently integrating novel episodes into existing frameworks of prior experience^1,3–6^. Beyond contextual discrimination, DG activity modulates downstream CA3 firing patterns, a network coordination essential for spatial working memory^7^.

The functional exclusion of the vast majority of DGCs from active ensembles is maintained by a combination of low intrinsic neuronal excitability, robust local GABAergic inhibition, and network oscillation-locked synaptic inputs^8–10^. Conversely, the subpopulation of DGCs recruited during environmental exploration exhibits higher basal excitability relative to the rest of the population^11^. Elucidating the mechanisms that govern neuronal eligibility and the active-to-silent ratio during encoding is critical, as functional disruption and abnormal reduction of ensemble size in aging, Alzheimer’s disease, and corresponding animal models is closely linked to impaired hippocampal coding and cognitive deficits^12–14^.

DGCs activated by experiences and recruited into neuronal ensembles are typically identified by real-time increases in *in vivo* calcium flux or the expression of immediate early gene (IEG) products such as FOS^1,6,15,16^. Following activation in a novel environment (NE), FOS^+^ DGCs differentially express numerous genes associated with broad functional categories, including transcription and signaling^16^. Despite its critical role in encoding and memory, the mechanisms that pre-configure a specific, small subpopulation of DGCs for ensemble eligibility remain elusive.

LINE-1 (long interspersed nuclear element 1, L1) retrotransposons comprise approximately 17% of the human genome and 19% of the mouse genome^17,18^. Although most genomic L1 copies are mutated, promoter truncated, or silenced by both DNA methylation and repressive histone modifications to maintain genomic stability^19^, certain evolutionarily “young” autonomous L1 elements are transcriptionally active in both the rodent and human brain^20,21^. While some L1 transcripts encode ORF1p and ORF2p to form ribonucleoprotein complexes capable of retrotransposition, most lack coding potential and exhibit only sporadic transposition in mature postmitotic neurons^20,22^. Emerging evidence demonstrates that untranslated L1 RNAs can act similarly to long noncoding RNAs to regulate neuronal differentiation^23–26^; however, L1 RNA is also expressed in mature neurons^20^, suggesting functions extending beyond development.

We confirmed high autonomous L1 RNA expression in mouse hippocampal neurons, including DGCs, but discovered that NE activated FOS^+^ DGCs express significantly lower L1 RNA levels than the rest of the neuronal population in the DG. Knocking down (KD) L1 RNA in the DG increased the number of NE-activated FOS^+^ DGCs and enhanced spatial working memory. Conversely, L1 RNA KD impaired pattern separation, a deficit aligning with an expanded ensemble size and denser coding, Overall, our study identifies L1 RNA as a vital negative regulator of DGC ensemble eligibility that supports sparse coding and preserves pattern separation by gating the size of the activated neuronal population.

## Results

### Differential DNA methylation between FOS^+^ and FOS^-^DGCs in single copy genes and transposons

Exposure to NE induces expression of the FOS protein, a proxy for neuronal activation, in a sparse population of DGCs in the mouse DG (**Fig. 1A**). Transcriptomes of FOS^+^ and FOS^-^ DGCs differ in a large number of genes, as previously reported by Lacar et al.^16^. Given that gene expression is immensely controlled by epigenetic mechanisms, including DNA methylation, we tested whether transcriptional differences between FOS^+^ and FOS^-^ DGCs have an underlying epigenetic basis.

**Fig. 1.**
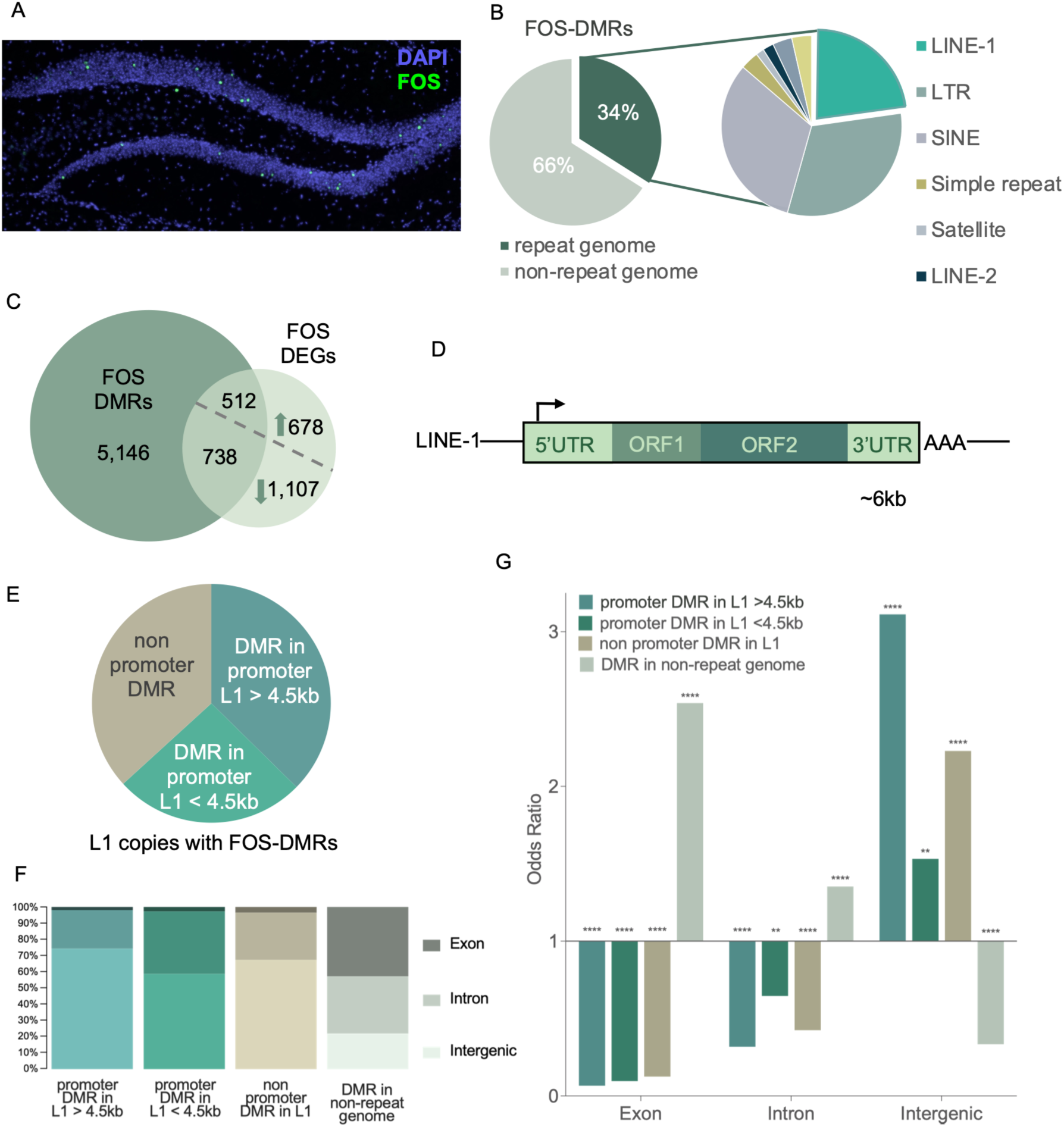
Differences in DNA methylation between NE activated FOS^+^ and non-activated FOS^-^ DGCs. (**A**) Activation of a sparse population of DGCs by NE (15 min) in the DG, visualized by immunofluorescence staining (IF) for FOS, 60 min after exposure, with DAPI counterstaining. (**B**) Distribution of FOS-DMRs in the non-repeat and repeat mouse genome, including various retrotransposons. (**C**) Significant overlap between FOS^+^ vs. FOS^-^ DMGs and DEGs (hypergeometric test p=2.876e-12, M=17925, K=6396, N=3034). (**D**) Structure of FL L1s with the transcription initiation site for *L1* RNA expression highlighted. (**E**) L1 DMRs are overrepresented in the 5’UTR/promoter sequence of L1 copies. Proportion of L1 DMRs in the promoter of FL and long (≥4.5 kb) copies and short (<4. 5kb) L1 copies. (**F**) Proportion of FOS-DMRs in genomic features. L1 promoter DMRs are predominantly intergenic while DMRs in the non-repeat genome are more evenly shared between exons, introns, and intergenic regions. (**G**) Distribution of FOS-DMRs across gene features as compared to randomized control regions. L1 promoter DMRs are predominantly intergenic while DMRs in the non-repeat genome are enriched in genic regions (exons and introns), Fisher’s exact test, ****p<0.0001, **p<0.01.

The DG was dissected 1h after NE exposure (NE), and FOS^+^ and FOS^-^ DGC nuclei were isolated by fluorescence activated nuclei sorting (FANS) using antibodies against FOS and the DGC marker PROX1. Sorted nuclei were then processed for enhanced reduced representational bisulfite sequencing (eRRBS)^27^. Based on our previous single nucleus RNA-sequencing data^28^, 88-90% of sorted PROX1+ DG nuclei exhibit a DGC specific transcriptome profile. We identified 30,016 differentially methylated sites (DMSs) between FOS^+^ and FOS^-^ nuclei (≥20% methylation difference, q<0.01, **Supplementary Table 1**). The majority of these sites clustered into 10,969 differentially methylated regions (DMRs) defined as having at least 2 DMSs within 1 kb (**Supplementary Table 2**). Among these, 5190 DMRs contained a mix of both hypo- and hypermethylated DMSs, while the remaining region was roughly evenly split between purely hypermethylated (2999) and purely hypomethylated (2780) DMRs.

Two-thirds of DMRs mapped to unique genic and intergenic regions. Notably, the remaining 34% (3,705 DMRs) mapped to repeat sequences (Repbase), predominantly to retrotransposons (L1, LTR, SINE), elements that are typically fully methylated and resistant to epigenetic remodeling in most somatic cells (**Fig. 1B, Supplementary Table 3**). The actual number of transposon associated DMRs is likely higher, as reads mapping to multiple loci were excluded from this analysis. Collectively, these data demonstrate that NE-activated DGCs differ from the resting population not only in the expression of a large number of genes^16^ but also in genome-wide DNA methylation.

Based on their ChromHMM chromatin states in DGCs^29^, FOS-DMRs within the unique, non-repetitive genome were significantly enriched in regulatory elements (both enhancers and promoters) and depleted in heterochromatic regions (**Extended Data 1A**). Differentially methylated genes (DMGs) that either contain DMRs or are in a putative regulatory relationship with DMRs^30^, showed a highly significant overlap with previously published NE FOS-DEGs^16^ (**Fig. 1C**, hypergeometric test p=2.876e-12). A similar proportion of these DMG-overlapping DEGs were upregulated and downregulated (43% and 40%, respectively). Over-Representation Analysis (ORA, webgestalt.org) revealed that the 512 upregulated DMGs (**Supplementary Table 4**) were enriched in similar Gene Ontology (GO) categories as the complete set of 1,190 upregulated DEGs (e.g., protein kinase/phosphotransferase activity and transcription factor binding, **Extended Data 1B, C**). Notably, these DEG x DMGs generally displayed a higher enrichment ratio than DEGs, suggesting that differential DNA methylation may directly contribute to the upregulation of these specific functional gene sets in FOS^+^ relative to FOS^-^ DGCs. Downregulated DEGs and DEG x DMGs showed no significant GO category enrichment, indicating that downregulated gene sets are not clustered around specific biological pathways.

Because transposons can exert a substantial influence on gene expression and cellular phenotype, we extended our study to the remaining one third of DMRs that mapped to repetitive elements (**Fig. 1B**). Since they can transcribe RNA, differential methylation of promoter containing full length (FL, ∼6 kb long) or near FL L1 copies (**Fig. 1D**) was particularly striking. Although typically lack coding potential due to acquired mutations, multiple studies have shown that these transcripts regulate gene expression independently of transposition^23–26^. Notably, thousands of FL and near-FL L1 copies exist in the mouse genome (and over a hundred in the human)^31^.

Intriguingly, most L1-associated-DMRs (64%) mapped directly to L1 promoter regions. These promoter-associated DMRs were identified not only in FL and relatively long (>4,5 kb) L1 copies, but also within shorter (<4,5 kb) truncated L1 elements (**Fig. 1E**). This strong concentration of DMRs within L1 promoters was highly unexpected because promoter-containing copies represent a minority of L1 insertions due to 5’ truncations (typically upstream of ORF2) in approximately 95% of L1 elements across L1 families^32^. We found that promoter-DMR containing FL and relatively long >4.5kb L1 copies primarily belonged to evolutionary young L1 families (L1MdA, L1MdGf, and L1MdTf). In contrast, shorter (<4.5 kb) L1 copies corresponded to more ancient evolutionary families (L1MdF, Lx, and L1N)^33^, consistent with the progressive accumulation of deletions and rearrangements over evolutionary time (**Supplementary Table 5**). Finally, L1 promoter DMRs were located predominantly within intergenic regions and were enriched in intergenic and depleted in genic regions, while non-repeat DMRs were located and enriched in exons and introns (**Fig. 1F, G**). Collectively, these findings demonstrate that a) methylation of L1 promoters exhibits a remarkable malleability in DGCs, b) the distinct FOS^+^ and FOS^-^ states are associated with differentially methylated L1 promoters, and c) L1 copies harboring promoter DMRs are typically physically segregated from single copy genes in the genome, suggesting a degree of transcriptional or functional autonomy.

### Reduced expression and altered nuclear distribution of *L1* RNA in activated FOS^+^ DGCs

Methylation of L1 promoter DMSs varied between 25-30% and 100% in non-activated FOS^-^ DGCs indicating substantial cell-to-cell variability in baseline methylation across hundreds of mapped L1 promoters in the DG (**Extended Data 2A**). Consistent with this bistable methylation pattern and its tilt toward the methylated state, L1 promoter DMRs were primarily associated with repressive histone marks and less prominently with active marks (**Extended Data 2B**). This pattern suggests that while most L1 promoter copies are silenced redundantly by CpG methylation and repressive chromatin modifications, a subset of unmethylated promoter copies residing within open chromatin remains active and drives L1 RNA transcription. Indeed, RNA fluorescent in situ hybridization (RNA FISH) using an ORF1 probe conserved among evolutionary young L1 elements revealed robust L1 RNA expression in the DG (**Fig. 2A**). This L1 RNA signal was almost exclusively nuclear, aligning with its nuclear retention and chromatin association reported in neurons^34^. Further, L1 promoter DMSs that fell within the lower baseline methylation range (25–60%) in FOS^-^ cells tended to show increased methylation in activated FOS^+^ DGCs, whereas those in the higher baseline range (90–100%) exhibited decreased methylation (**Extended Data 2A**).

**Fig. 2.**
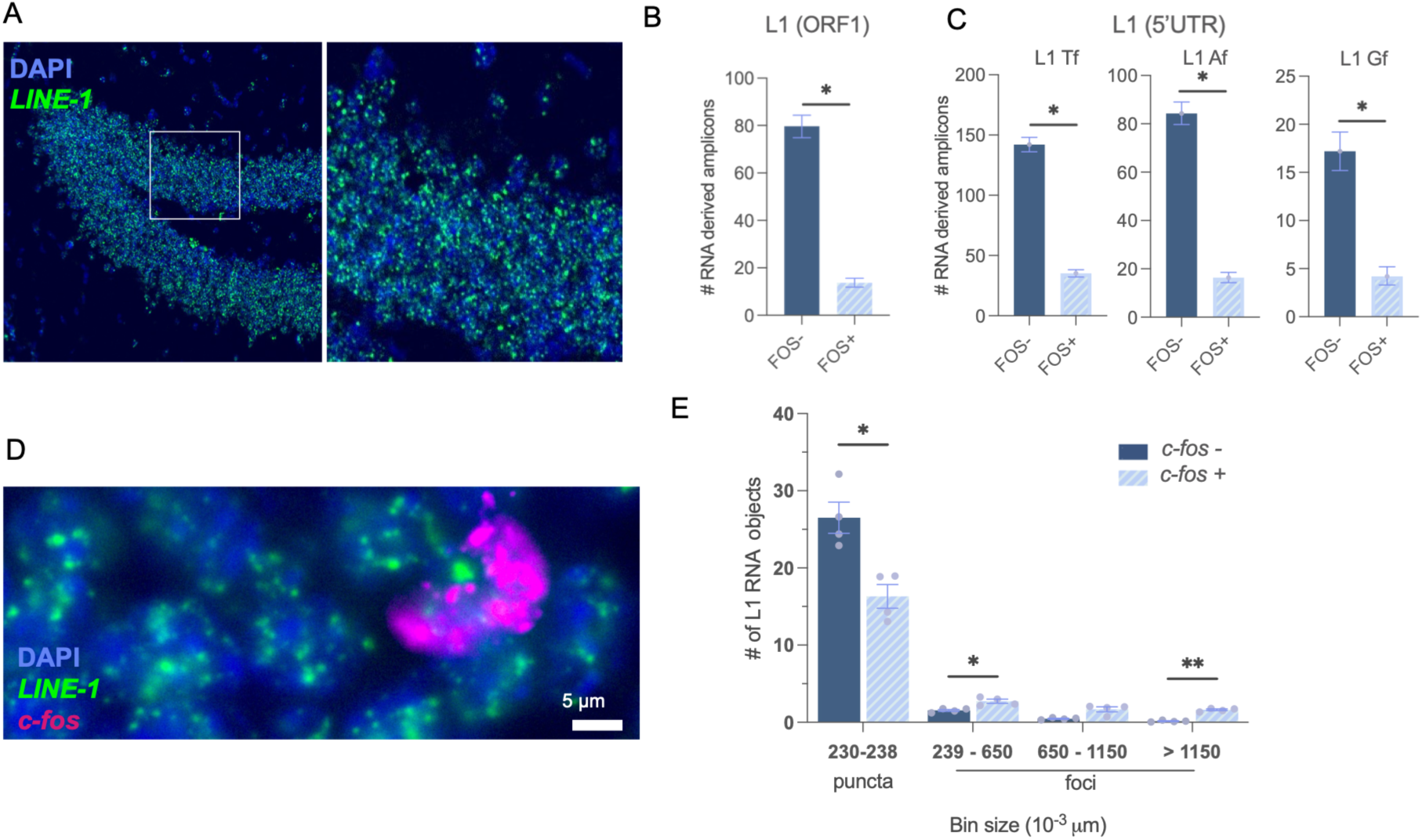
Reduced expression of L1 RNA in activated FOS^+^, relative to non-activated FOS^-^DGCs. (**A**) Expression of L1 RNA in the dorsal DG. FISH with an L1 ORF1 probe conserved across all young family members. Magnified image shows that L1 RNA is nuclear. (**B**) dRT-PCR assay with an ORF1 probe set specific for young L1 families shows reduced number of L1 RNA-derived amplicons in FOS^+^ relative to FOS^-^ DGCs, standard deviation of the confidence intervals difference (see Methods), *P<0.05. Error bars indicate the Poisson 95% confidence intervals. (**C**) Same as B but with young family specific probe sets, *P<0.05. (**D**) Representative images of L1 RNA in puncta in a c-*fos*^-^ and in foci in a c-*fos*^+^ DGC. FISH with a conserved ORF1 probe, 30 min after environmental exposure. (**E**) Histogram of size distribution of the L1 RNA FISH signal in FOS^+^ and FOS^-^ cells. Two-way ANOVA, main effect of cell type, F (1,3) = 20.52, P = 0.021; main effect of size F (1,016,3.04) = 806.50, P = 0.001. Bonferroni’s multiple comparisons tests *P<0.05, ** P<0.005. N = 4 animals per group.

Given the established inverse correlation between promoter methylation and transcription, shifts in L1 promoter methylation could modulate L1 RNA abundance in FOS^+^ relative to FOS^-^ DGCs^35^. However, considering the epigenetic redundancy involved in retrotransposon silencing^36^, the localized hypomethylation of certain L1 copies alone might not automatically trigger transcription. In contrast, locus specific hypermethylation in FOS^+^ DGCs is expected to silence baseline expression. This scenario predicts a net overall reduction in L1 RNA transcripts within activated neurons. Supporting this hypothesis, total and family-specific L1 RNA levels were significantly reduced in FANS-isolated FOS^+^ nuclei relative to FOS^-^ DGC nuclei (**Fig. 2B, C**).

Beyond total abundance, the spatial localization of L1 RNA differed dramatically between activated *c-fos^+^*/ FOS^+^ and non-activated *c-fos^-^/* FOS^-^ DGCs. In resting *c-fos^-^*DGCs, the L1 RNA signal appeared predominantly as diffuse puncta with only a few larger foci. In contrast, activated *c-fos^+^* DGCs exhibited fewer puncta and a marked accumulation of large, distinct foci (**Fig. 2D-E**). Notably, these large L1 RNA FISH foci in *c-fos^+^* cells overlapped with the immunofluorescence signal of ZCCHC8, a core structural scaffold of the nuclear exosome targeting (NEXT) complex^37^. This overlap was weaker for PABPN1, a key adapter molecule for the poly(A) tail exosome targeting (PAXT) complex^38^ (**Extended Data 3A, B**). Together, these spatial and biochemical dynamics suggest that the reduction of diffuse L1 RNA puncta and the concurrent appearance of L1 nuclear foci in activated *c-fos^+^* DGCs are driven by an enhanced recruitment of L1 transcripts to specialized nuclear exosome complexes responsible for nuclear RNA surveillance and degradation.

### Antisense knockdown of *L1* RNA in the dentate gyrus increases the number of DGCs activated in a novel environment

To test whether low L1 RNA abundance in FOS^+^ DGCs is a cause or a consequence of neuronal activation, we downregulated L1 transcripts using a previously validated, nuclease-resistant L1 antisense oligonucleotide^39^ (ASO), delivered by bilateral microinjection into the dorsal DG (**Fig. 3A**). The ASO sequence corresponded to the conserved ORF1 region of young L1 elements and contained no other complementary sequences within the mouse reference genome (GRCm39/mm39). Given its position near the 5’ end of the L1 transcript, this ASO preferentially targeted RNA derived from autonomous L1 copies. Both Cy3 ASO and scrambled control oligo (ScrO) reached the nuclear compartment of DGCs, as evidenced by robust nuclear Cy3 fluorescence at day three post-injection (**Fig. 3B**). Quantitative flow cytometry confirmed that virtually all Prox1+ nuclei isolated from dissected DGs were Cy3-positive and that these nuclei had comparable Cy3 signal intensity, reflecting a uniform cellular uptake of the oligonucleotides across the DGC population (**Fig. 3C**, P4 gating, and **Extended Data 4**). To verify KD, we performed digital RT-PCR (dRT-PCR) on RNA isolated from FANS-sorted FOS^+^ and FOS^-^ nuclei. ASO treatment significantly reduced the number of L1 RNA-derived amplicons compared to ScrO-infusion (**Fig. 3D**). Because the targeted sequence was highly conserved, the ASO successfully downregulated L1 RNA across all three major active young L1 families (L1MdA, L1MdGf, and L1MdTf) (**Fig. 3E**). Furthermore, ASO delivery successfully reduced the number of L1 RNA puncta in FOS^-^ DGCs down to levels observed in FOS^+^ DGCs treated by control ScrO (**Fig. 3F**). ASO administration failed to decrease the number of L1 RNA foci (**Fig. 3G**), suggesting that transcripts sequestered within these macromolecular nuclear complexes are physically shielded or otherwise inaccessible to antisense-mediated degradation.

**Fig. 3.**
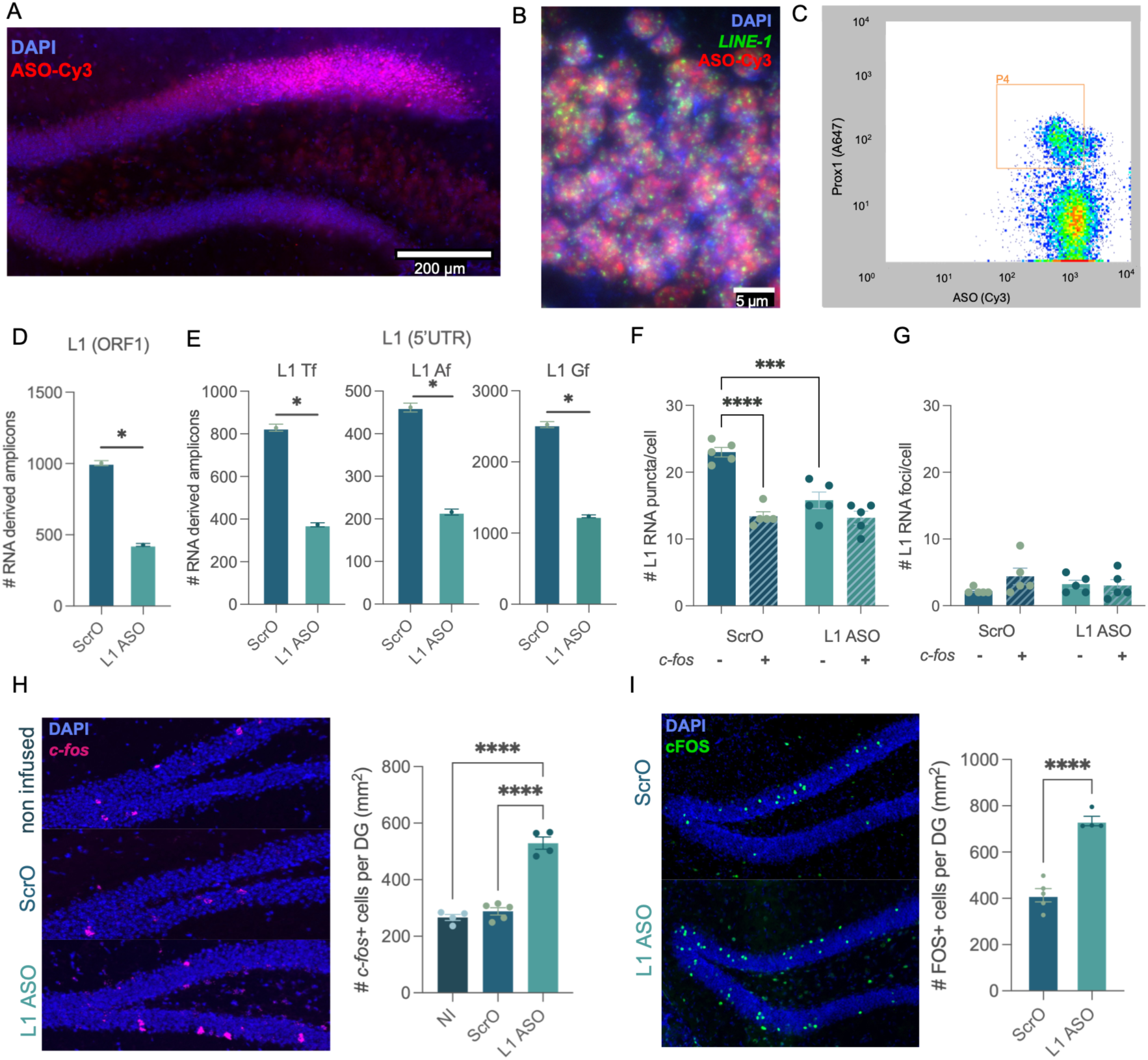
ASO mediated knockdown of L1 RNA increases the number of activated, *c-fos^+^*/FOS^+^, cells in the DG following novel environment exposure. (**A**) Cy3 labeled FANA (2′-deoxy-2′-fluoro-D-arabinonucleic acid) ASO, injected close to the upper blade of the granule cell layer one week after injection. (B) High magnification of DGCs with DAPI counter-staining showing the nuclear localization of Cy3 ASO. (C) Flow cytometry shows that virtually all PROX1+ (as well as PROX1-) DG nuclei took up Cy3 ASO. (**D**) ASO downregulated *L1* RNA levels in FANS DGC nuclei 3 days after delivery. dRT-PCR assay with an ORF1 probe set specific for young L1 families, standard deviation of the confidence intervals difference, *P<0.05. Error bars indicate the Poisson 95% confidence intervals. (**E**) Same as D but with family specific probe sets, *P<0.05. (**F**) ASO reduced the number of *L1* RNA puncta in c-fos^-^ DGCs to a level measured in ScrO *c-fos*^+^ DGCs. Two-way ANOVA, Main effect of treatment, F (1,16) = 15.92, P=0.001; main effect of activation state, F (1,16) = 43.27, P<0.0001, interaction, F (1,16) = 14.24, P=0.002. Tukey’s multiple comparison test, ***P=0.0003, ****P<0.0001. N=5 ScrO, N=5 ASO. (**G**) ASO had no significant effect on the number of L1 RNA foci. Main effect of treatment, F (1,16) = 0.06, P=0.812; main effect of activation state, F (1,16) = 1.46, P=0.244. (**H**) ASO increased the number of *c-fos*^+^ DGCs activated in a NE, 3 days post-infusion. Left: Representative images of *c-fos^+^*cells in ASO, ScrO, and non-infused (NI) DGs, 30 min after NE exposure. Right: Quantification of *c-fos*^+^ ensemble expansion by ASO. One-way ANOVA, F (2,10) = 84.34, P<0.0001. Tukey, ****P<0.0001. N=4 NI, N=5 ScrO, N=4 ASO. (**I**) ASO increased the number of FOS^+^ DGCs activated in NE, 7 days post-infusion. Left: Representative images, 1h after NE exposure. Right: Quantification of FOS^+^ ensemble expansion by ASO. Unpaired two-tailed t-test: t_7_=8.66, ****P<0.0001.

Next, we evaluated whether ASO-induced depletion of L1 RNA in the DG alters NE-induced neural activation. Remarkably, three days after oligonucleotide delivery, ASO treatment nearly doubled the size of the neural ensemble activated by novel environment (NE) exposure compared to non-infused and ScrO control groups, as measured by *c-fos* RNA FISH (**Fig. 3H**). This expansion of the activated population persisted up to 7 days post-delivery, as verified by FOS immunohistochemistry (IHC, **Fig. 3I**) indicating a long-lasting effect of ASO on NE-induced ensemble size. While the number of FOS^+^ DGCs was predictably lower in home-cage controls, ASO infusion still led to a significant expansion of the home-cage FOS^+^ ensemble, approaching the size of the NE-induced ensemble in control ScrO mice (**Extended Data 5**). Consequently, ASO-induced ensemble expansion is not restricted to novelty-driven neuronal activation, but rather represents a generalized phenomenon in the DG that occurs independently of whether activation is induced by robust novel stimuli or subtle ambient home-cage signals. Taken together, these data indicate that lowering L1 RNA levels acts as a switch permitting the transition of DGCs into the ensemble-eligible state.

Neural stem cells in the subgranular zone (SGZ) of the adult DG proliferate and continuously produce young neurons, which exhibit higher baseline excitability than mature DGCs. However, ASO delivery influenced neither the number of Ki67+ proliferating cells in the SGZ 3 days post-infusion nor the number of DCX positive immature neurons in the SGZ and the entire DG 7 days post-infusion (**Extended Data 6A, B**). These data indicate that primarily mature neurons contribute to the expansion of NE activated ensemble in ASO mice. This conclusion is consistent with the relatively even distribution of ASO induced FOS^+^ cells across the granule cell layer, as well as the documented low recruitment rate of immature DGCs into behaviorally activated ensembles^40^. Furthermore, the expansion of the active ensemble occurred within 3 days of ASO infusion, representing a significantly faster temporal dynamic than the functional integration of adult-born neurons, a developmental process that requires at least 1–3 weeks^41^. Together, these findings demonstrate that L1 RNA KD acts directly on mature DGCs to increase their ensemble eligibility, independent of adult neurogenesis.

### Antisense knockdown of L1 RNA increases excitability in DGCs

Increased relative excitability and functional connectivity have been reported to bias hippocampal projection neurons toward the ensemble eligible state and facilitate their allocation to a coding ensemble^42^. Because L1 ASO delivery expanded the active neuronal ensemble, we tested whether the downregulation of L1 RNA levels in the DG alters DGC excitability and synaptic communication with upstream circuit partners. In whole-cell current-clamp recordings of individual DGCs^1^ during slice electrophysiology recordings^43^ (**Fig. 4A**), we found that the resting membrane potential (RMP) was similarly near -70 mV in both ASO and ScrO groups (**Fig. 4B**). However, ASO-treated DGCs exhibited an enhanced capacity for action potential (AP) firing across increasing step injections of depolarizing current (**Fig. 4C-D**). This increase in repetitive firing occurred in the absence of significant changes in the threshold or rheobase for AP firing (**Extended Data 7A, B**). Passive and active membrane properties and measures of ion channel conductances contributing to intrinsic excitability were also similar between groups, including membrane resistance, a measure of overall channel conductance/leak (**Extended Data 7C**). Furthermore, the kinetics of the first evoked AP, including the rise slope, drop slope, and afterhyperpolarization potential (AHP) magnitude were unchanged between groups, indicating intact voltage-gated Na+ and K+ channel dynamics in ASO infused cells (**Extended Data 7D-F**). Voltage sag, a measure of HCN channel-mediated h current elicited in response to hyperpolarizing current injection, was similarly unaffected (**Extended Data 7G**).

**Fig. 4.**
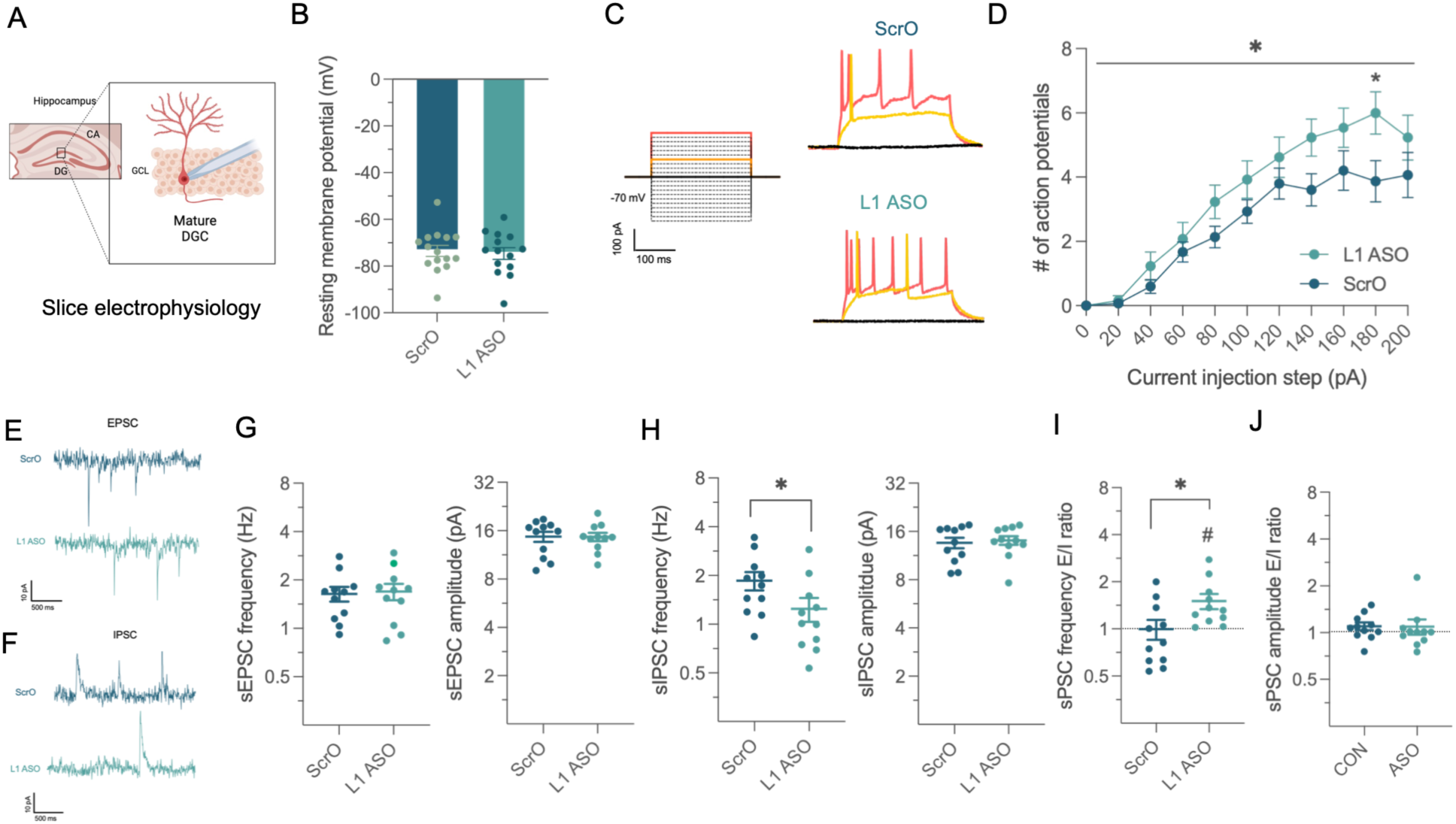
ASO increases the excitability of DGCs and decreases synaptic inhibition. **A.** Schematic of whole-cell slice electrophysiology recordings of DGCs in the DG. **B.** The resting membrane potential (RMP) of ScrO DGCs (n=15 cells/4 mice) and ASO (n=14 cells/4 mice) was similar (unpaired two-tailed t-test: t_27_=0.312, P=0.757). **C.** Stimulus waveform for current-injected action potentials, from a holding potential of −70 mV, consisting of a series of depolarizing current steps ranging from −200 to 300 pA in 20 pA increments (left). Representative traces of action potentials fired in response to 0 pA, 80 pA and 200 pA steps of current injection (right). **D**. DGCs from ASO mice fired more APs in response to increasing 20 pA steps of current from a common holding potential of -70 mV (15 cells/4 ScrO mice and n=13 cells/4 ASO mice). Two-way RM ANOVA: main effect of current step F (10,250) = 68.63, P<0.0001, and treatment x current step interaction F (10,260) = 2.00, *P=0.034; with post hoc t-tests with Holm-Sidak corrections for multiple comparisons between ScrO and ASO at each step: *P=0.023. **E.** Representative traces of sEPSCs in ScrO (top) and ASO (bottom) DGCs. **F.** Representative traces of sIPSCs in ScrO (top) and ASO (bottom) DGCs. **G.** sEPSC frequency (left) and amplitude (right) in DGCs were not altered by L1 ASO KD (P>0.05) n=11 cells/4 ScrO mice and n=11 cells/4 ASO mice. **H.** sIPSC frequency (left; t_20_=2.24, ^∗^P=0.037), but not amplitude (right: P>0.05), was lower in ASO than ScrO DGCs. **I.** The excitatory/inhibitory (E/I) ratio for sPSC frequency was higher in ASO than ScrO DGCs (unpaired t-test, t_10_=2.76, ^∗^P=0.012) and significantly shifted above 1.0 toward excitation in ASO cells (one-sample t-test: t_10_=3.04, ^#^P=0.013) but not ScrO cells (one-sample t-test: t_10_=0.03, P=0.974). **J.** The E/I ratio for sPSC amplitude was neither different between ScrO and ASO DGCs (P>0.05) nor significantly different from 1.0 in either group (P>0.05). All data are shown as mean ± SEM with individual data points represented when possible.

To isolate spontaneous excitatory and inhibitory postsynaptic currents (sEPSCs and sIPSCs, respectively) within individual DGCs, we also performed voltage-clamp recordings from a holding potential of -55 mV and +10 mV (**Fig. 4E, F**). The frequency and amplitude of sEPSCs were similar between ScrO and ASO conditions (**Fig. 4G**), indicating that glutamatergic synaptic transmission remained unaltered. While sIPSC amplitudes were also similar between groups, the frequency of sIPSCs was significantly lower in ASO-treated DGCs compared to ScrO controls (**Fig. 4H**), suggesting a reduction in synaptic inhibition. Consequently, this selective loss in synaptic inhibition drove a significant shift toward excitation in the excitatory/inhibitory (E/I) balance of sPSC frequency in ASO infused cells that was significantly higher than that in ScrO cells, with no difference in E/I balance for amplitude (**Fig. 4I, J**). Together, these results suggest that reduced basal synaptic inhibition of ASO DGCs may contribute to their ability to fire more upon robust depolarization.

### Antisense knockdown of *L1* RNA improves spatial working memory and modulates pattern separation

The dorsal DG has been implicated in supporting multiple distinct mnemonic functions, with coordinated network activity within this structure promoting spatial working memory and goal-directed behavior^7^. Mice with bilateral ASO infusion into the dorsal DG (cohorts 1 and 3, **Fig. 5A**) exhibited enhanced performance in the continuous spontaneous alternation task in the Y maze relative to both ScrO-infused and non-infused control groups (**Fig. 5B**). To evaluate working memory under higher cognitive demand, we tested mice in a discrete T-maze alternation task with a 30-second inter-trial delay, a paradigm that requires holding spatial representations internally in the absence of ongoing sensory input^44^. ASO-treated mice displayed better performance compared to controls in this challenging task (**Fig. 5C**, cohorts 2 and 3). The enhanced spatial working memory of ASO-treated mice was independent of general ambulation, as total locomotion was indistinguishable among the ASO, ScrO, and non-infused groups (**Fig. 5D**). Taken together, these data demonstrate that ASO mice with an enlarged DG ensemble exhibit increased spatial working memory.

**Fig. 5.**
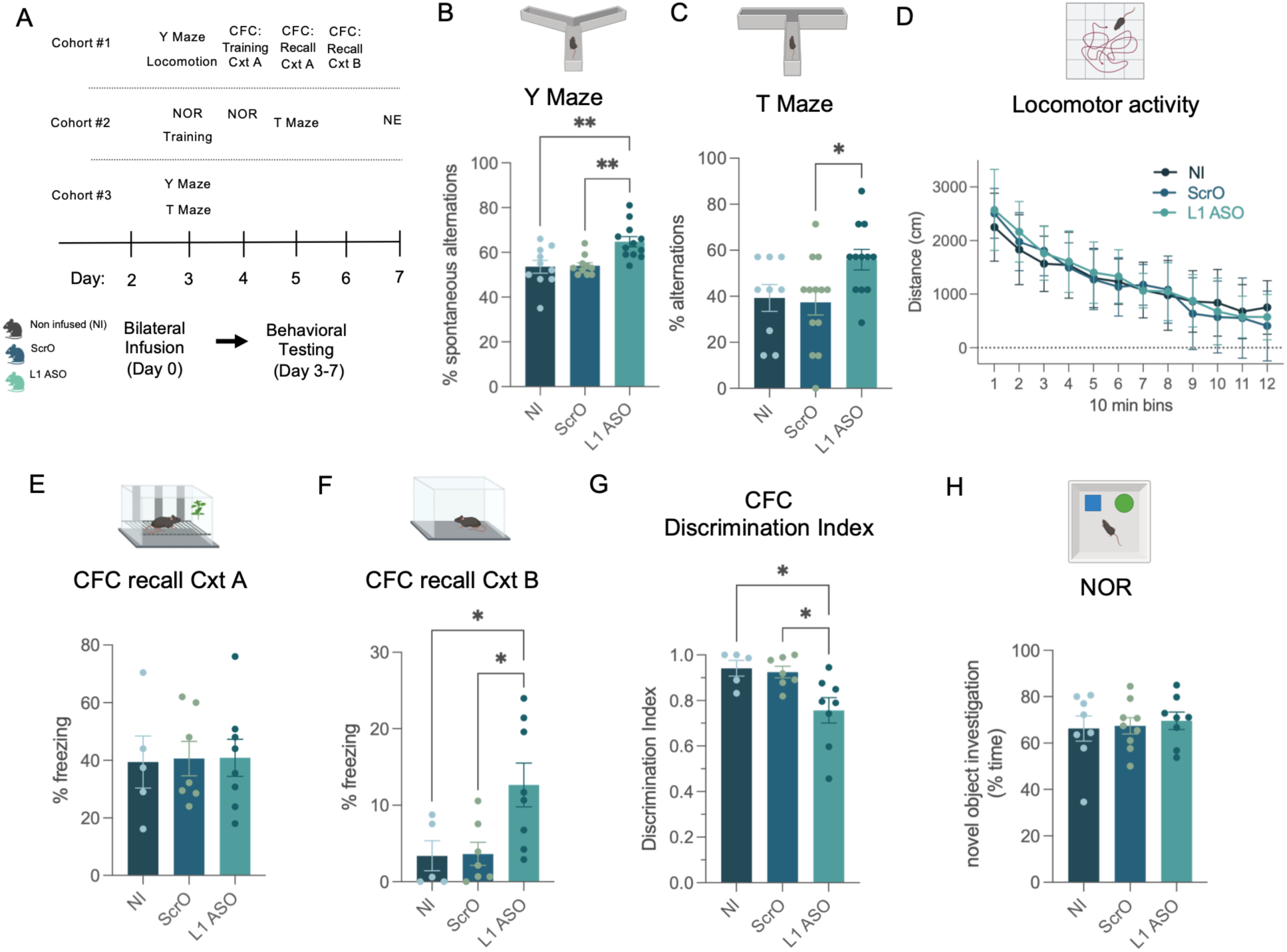
Behavioral testing of ASO injected mice in comparison with ScrO injected and non-infused (NI) controls in hippocampal dependent and independent behaviors. (**A**) Timeline of behavioral testing in three cohorts of animals after ASO and ScrO delivery. (**B**) Spontaneous alternations in the Y maze (cohorts 1 and 3). One-way ANOVA, F (2,29) = 8.17, P=0.001, Tukey multiple comparisons **P<0.01, N=12 (ASO), 10 (ScrO), 10 (NI). (**C**) Spatial working memory in delayed non-matching to place (DNMTP) task in T maze with 30 sec delay (cohorts 2 and 3). One-way ANOVA, F (2,31) = 3.93, P=0.03, Tukey multiple comparisons *P=0.035, N=12 (ASO), 13 (ScrO), 9 (NI). (**D**) 120 min locomotor activity (cohort 1) in 10 min bins, N=7 (ASO), 6 (ScrO), 11 (NI). (**E**) CFC recall in context A, ANOVA F (2,17) = 0.01, P=0.989, N=8 (ASO), 7 (ScrO), 5 (NI). (**F**) Same as E but in context B, F (2,17) = 5.37, P=0.016, Tukey *P<0.05. (**G**) Discrimination index, reflecting discrimination between contexts A and B, is lower in ASO than in control mice. One-way ANOVA, F (2,17) = 5.53, P=0.014, Tukey *P<0.05. (**H**) ASO treatment had no effect on performance in the non-hippocampal novel object recognition test. One-way ANOVA, F (2,22) = 0.15, P=0.860, N=8 (ASO), 9 (ScrO), 8 (NI).

Beyond working memory, a canonical function of the DG is pattern separation, a process by which overlapping entorhinal cortex inputs representing similar contexts are orthogonalized into distinct DGC assemblies^4,6,40^. This function is generally studied using the contextual fear conditioning (CFC) paradigm^40^. Following CFC, ASO-treated, ScrO-treated, and non-infused control mice exhibited comparable freezing behavior when reintroduced to the conditioned context (Context A) 24 hours later, indicating that L1 KD does not alter baseline associative fear memory acquisition or expression (**Fig. 5E**). However, exposure on the subsequent day to a novel, similar environment (Context B, featuring modified wall patterns and floor textures) elicited minimal freezing in ScrO and non-infused controls, but significantly more freezing behavior in ASO-treated mice (**Fig. 5F**). Elevated fear response suggests a deficit in context discrimination. This impairment was further substantiated by a significantly lower discrimination index in the ASO group relative to controls (Fig. 5G), providing a quantitative metric of the diminished capacity of ASO-treated mice to discriminate between similar environments (**Fig. 5G**). These behavioral findings align with the theoretical model that an artificial, ASO-induced enlargement of the active DGC ensemble disrupts the neuronal sparsity and orthogonal coding required to statistically segregate overlapping environmental contexts within the DG.

As anticipated, mice infused with ASO in the DG and control animals exhibited comparable performance in the non-hippocampal novel object recognition (NOR) test (**Fig. 5H**). The similar novelty preference across cohorts indicates that the enhanced working memory observed in ASO mice during Y- and T-maze testing was not driven by heightened novelty-seeking behavior. Furthermore, ASO KD within the dorsal DG did not alter anxiety-like behavior in the open field test. This suggests that the behavioral effects of the ASO are specific to spatial cognitive tasks, without influencing emotional behaviors traditionally modulated by the medial prefrontal cortex (mPFC) and the ventral hippocampus, including the ventral DG^45^ (**Extended Data 8**).

Following the conclusion of behavioral testing, NE exposure on day 7 of cohort 2 ASO mice consistently yielded an expanded FOS^+^ ensemble 1 hour post-exposure (**Fig. 3I**). These findings demonstrate that repeated behavioral testing does not confound or diminish the ASO’s impact on ensemble size. Collectively, these data establish L1 RNA as a critical regulator of DG ensemble size and DG dependent behaviors.

## Discussion

Sparse coding and restricted ensemble sizes within the cortex and hippocampus offer distinct computational advantages, including high efficiency and low metabolic costs for representing experiences^1,2^. While this coding scheme was initially found in rodents, it has since been validated by direct recordings in the human hippocampus^3^. Mechanistically, a relative increase in excitability, regulated by the transcription factor CREB (cAMP response element-binding protein), has been strongly implicated in preconfiguring neurons for allocation to memory engrams^11,42^. However, the upstream mechanisms governing excitability and ensemble eligibility within sparse neuronal populations remain elusive. Here, we identify a single molecular entity, RNA transcribed from autonomous L1 retrotransposons within the mouse genome, as a key regulator of sparse coding and restricted ensemble sizes in the DG.

Although the mouse genome harbors approximately 6,000 to 7,000 autonomous L1 elements, they are predominantly silenced through the redundant mechanisms of promoter methylation and heterochromatinization. Nevertheless, L1 RNA expression occurs in the rodent (and human) brain^20,21^, and we demonstrate here that young L1 family-specific transcripts are abundantly expressed in mouse DGCs. Consistent with this expression profile, a fraction of L1 promoters remains unmethylated and resides within an actively transcribing chromatin state. Furthermore, our data show that L1 RNA expression is dynamically regulated in DGCs; specifically, FOS^+^ cells activated by novel environment (NE) exposure exhibit significantly lower L1 RNA levels than the rest of the DGC population. While the precise regulatory mechanism remains to be fully elucidated, switching CGs at select L1 promoters from the unmethylated to the methylated state in FOS^+^ DGCs likely drives transcriptional silencing, thereby reducing L1 RNA copy numbers. Conversely, while a subset of L1 promoters exhibited hypomethylation in FOS^+^ DGCs, such epigenetic shifts require a concomitant loss of heterochromatin to permit promoter activation. We therefore hypothesize that L1 promoter methylation and transcriptional repression predominate over activation in FOS^+^ cells, resulting in a net downregulation of L1 RNA in a subset of DGCs primed for activation. Given that activated ensembles maintain a remarkably consistent size across diverse behavioral modalities, we propose a stochastic hypermethylation mechanism that transiently reduces L1 RNA levels within a sparse population of DGCs, thereby establishing a permissive molecular threshold required for ensemble eligibility and activation at any given environment. As a non-mutually exclusive mechanism, accelerated nuclear L1 RNA degradation may also occur within a sparse population of DGCs, a hypothesis supported by the spatial aggregation of the L1 RNA signal and its co-expression with nuclear exosome complexes in activated DGCs.

Next, we tested whether artificially lowering L1 RNA levels across the DGC population would result in an ensemble eligible and activatable cell phenotype. Cells configured for ensemble recruitment exhibit a relative increase in excitability, and we show that ASO-mediated reduction of L1 RNA in DGCs attenuates GABAergic synaptic inhibition and elevates repetitive firing, without overtly altering the channel conductances that contribute to intrinsic excitability or early AP firing. We posit that during hippocampus-dependent learning, reduced basal inhibition alters the E/I balance that preserves sparse coding, driving increased DGC firing and thereby expanding the active coding ensemble in ASO-treated mice. Collectively, these findings indicate that L1 RNA downregulation promotes, and is likely required for, the transition of DGCs into a hyperexcitable, ensemble-eligible, and subsequently activated state. Given the orthogonal coding scheme and pattern separation of contexts in the DG, it follows that the subset of DGCs with low L1 RNA levels must be larger than the population actually activated in a given environment; however, the precise dimensions of this ensemble eligible pool remain to be determined. Taken together, our data establish a causal relationship between L1 RNA levels, ensemble eligibility, and activated ensemble size, identifying L1 RNA as a critical negative regulator of neuronal allocation to memory traces in the DG.

L1 RNA has previously been shown to control mouse and human embryonic stem cell identity and transcriptional programs during mouse and human early embryogenesis^24^, as well as neuronal differentiation during brain development^34,46^. Mechanistically, L1 RNA has been reported to modulate gene expression predominantly as a chromatin-associated transcript, recruiting epigenetic modifiers to repress or promote local gene expression depending on genomic context^24,39^. For instance, in early embryos, L1 RNA recruits Nucleolin and Kap1 to silence the transcription factor Dux and 2-cell specific genes, thereby driving exit from the totipotent state, while simultaneously binding ribosomal DNA to stimulate rRNA synthesis^24^. Paralleling these developmental roles, we hypothesize that the transcriptomic divergences observed between FOS^+^ and FOS^-^ DGCs are orchestrated by the differential abundance of L1 RNA within these distinct cellular populations.

Notably, L1 RNA functions have been independently co-opted in mice and humans through convergent evolution to sustain early embryonic development. The human genome contains approximately 100 autonomous L1 copies, most of which, like their murine counterparts, are retrotransposition-incompetent and do not encode functional proteins^31,47^. Given this evolutionary convergence and the documented expression of L1 RNA in human postmitotic neurons, it is plausible that L1 RNA similarly regulates ensemble eligibility and neuronal allocation in the human brain.

Consistent with an expanded ensemble size and denser network coding, ASO-mediated KD of L1 RNA in the DG enhanced spatial working memory in Y- and T-maze alternation tests, while concurrently disrupting pattern separation. Working memory and pattern separation represent core cognitive domains that are characteristically impaired in neurodegenerative diseases and other neurological pathologies^48–51^. Importantly, our findings carry profound therapeutic implications, as ASO-mediated downregulation of hippocampal L1 RNA may restore functional ensemble sizes, rescue working memory, and preserve spatial discrimination in pathological contexts characterized by functional and structural neuronal loss, such as neurodegeneration and stroke. For example, the DG is among the earliest and most vulnerable structures affected in Alzheimer’s disease (AD), and AD patients exhibit prominent deficits in working memory^51^ and spatial pattern separation^48–50^ from early clinical stages.

In summary, our work shows that L1 copies and their expressed RNA have been co-opted during evolution to govern sparse coding and pattern separation by restricting intrinsic excitability and ensemble eligibility to a small population of neurons in the DG. While prior studies identified elevated excitability as a hallmark of the ensemble-eligible state, our study is the first to identify a molecular master regulator of excitability, ensemble eligibility, and active ensemble size in the DG, and by extension, spatial working memory and spatial discrimination.

## Acknowledgments

We would like to thank the support of the Epigenomics Core and the Flow Cytometry Core Facility of Weill Cornell Medicine and Dr. Jihye Kim and Dr. Kenneth Wayne Johnson for their help in the behavioral experiments. We also thank AUM BioTech, LLC for providing AUM *silence^TM^*sdASO^TM^ and technical support. Figures were created in BioRender.

## Author contributions

Conceptualization: NP, KP, MT

Methodology: NP, PUH, IP, JPW, FT, HN, KP, MT

Investigation: NP, JGT, IP, JPW, HN, MT

Visualization: NP, PUH, IP, JPW, HN, RS,

Funding acquisition, KP, MT

Writing – original draft: MT

Writing – review & editing: NP, KP, MT

## Supplementary Materials

Extended Data, Figs. S1-S8

Supplementary Tables 1-5

