## Extended Data for "Expanding Hippocampal Ensembles through LINE-1 RNA Downregulation Enhances Cognition"

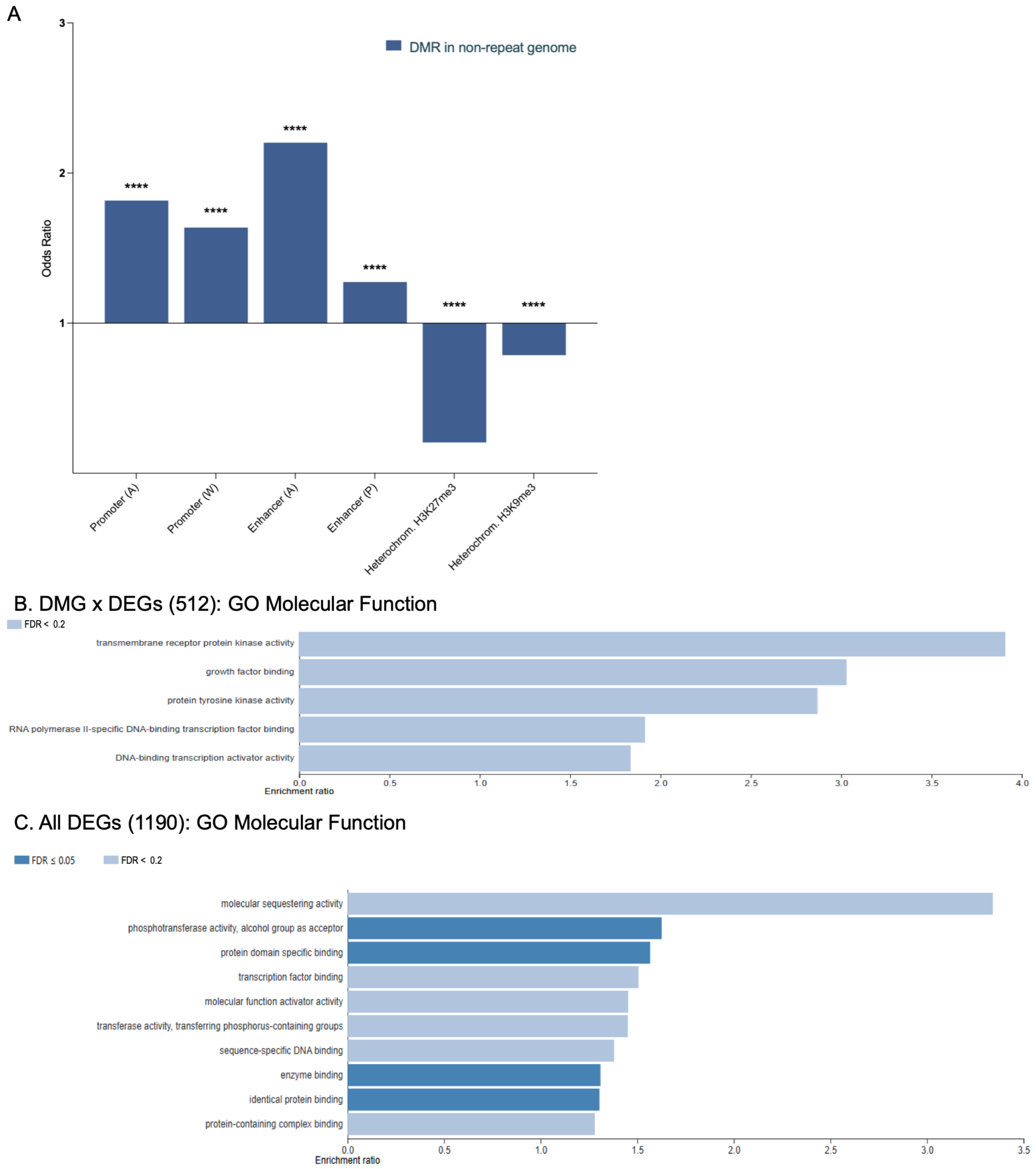


**Extended Data 1.**

**Genomic features of FOS-DMRs and their association with GO functional categories.** (**A**) DMRs in the non-repeat genome were enriched in enhancers and promoters, while depleted in heterochromatic regions. (**B, C**) Over-Representation Analysis (webgestalt.org) of the 512 FOS-DMG x DEGs (B) and the 1190 FOS-DEGs (C). Category size was between 5 and 1000, redundancy reduction was applied with weighted set cover, and the number of permutations was set to 1000.

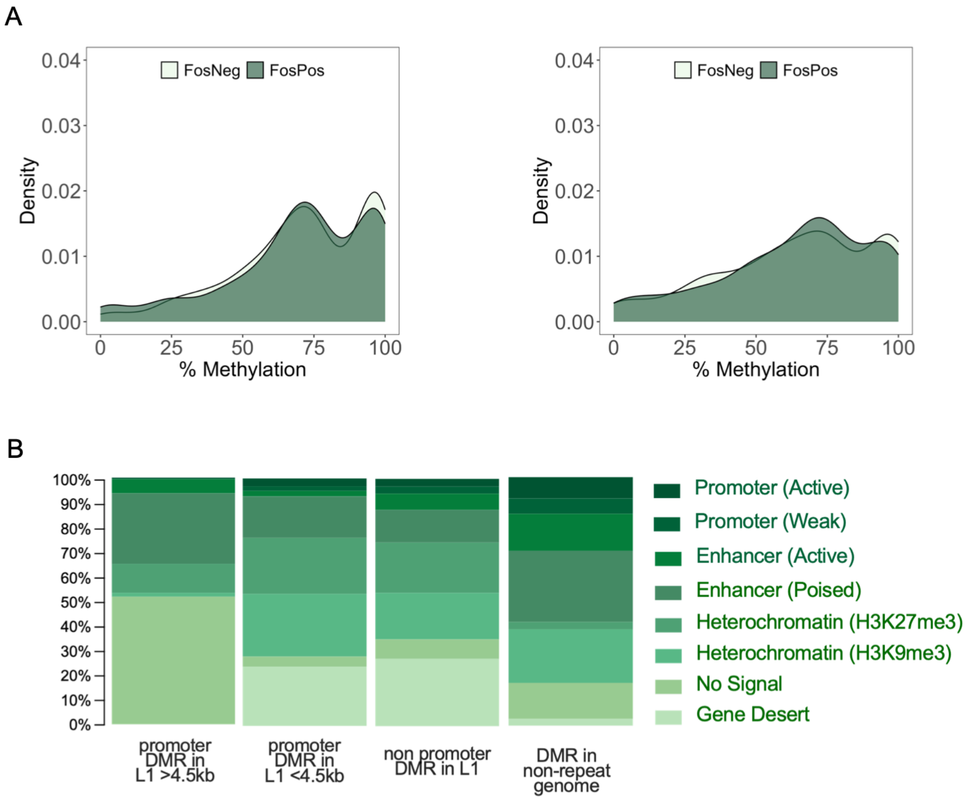


**Extended Data 2.**

**Epigenetic signatures of L1 promoter DMRs in DGCs. (A**) Methylation based distribution of FOS-DMSs within promoter DMRs in >4.5kb (left) and <4.5kb (right) L1 copies. (**B**) Chromatin states of L1 promoter DMRs and DMRs in non-promoter L1 copies, based on previous ChromHMM chromatin states in mouse DGCs. For comparison, the chromatin states of FOS-DMRs in the non-repeat genome are also shown.

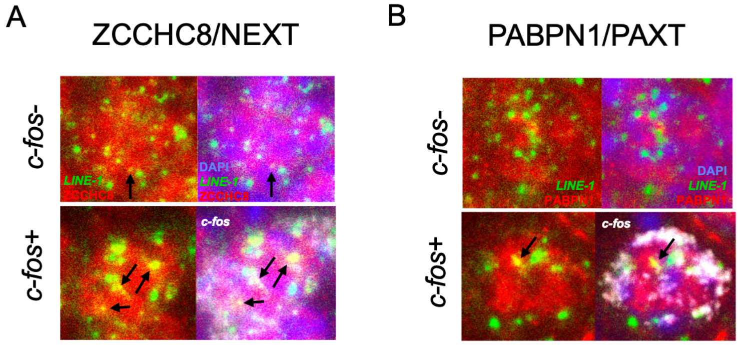


**Extended Data 3.**
**Overlap of L1 RNA in aggregates with ZCCHC8 (A) and PABPN1 (B) signal in *c-fos*+ cells, indicated by arrows**. FISH (L1 RNA, green) combined with IF (ZCCHC8 and PABPN1, red, and *c-fos*, white).

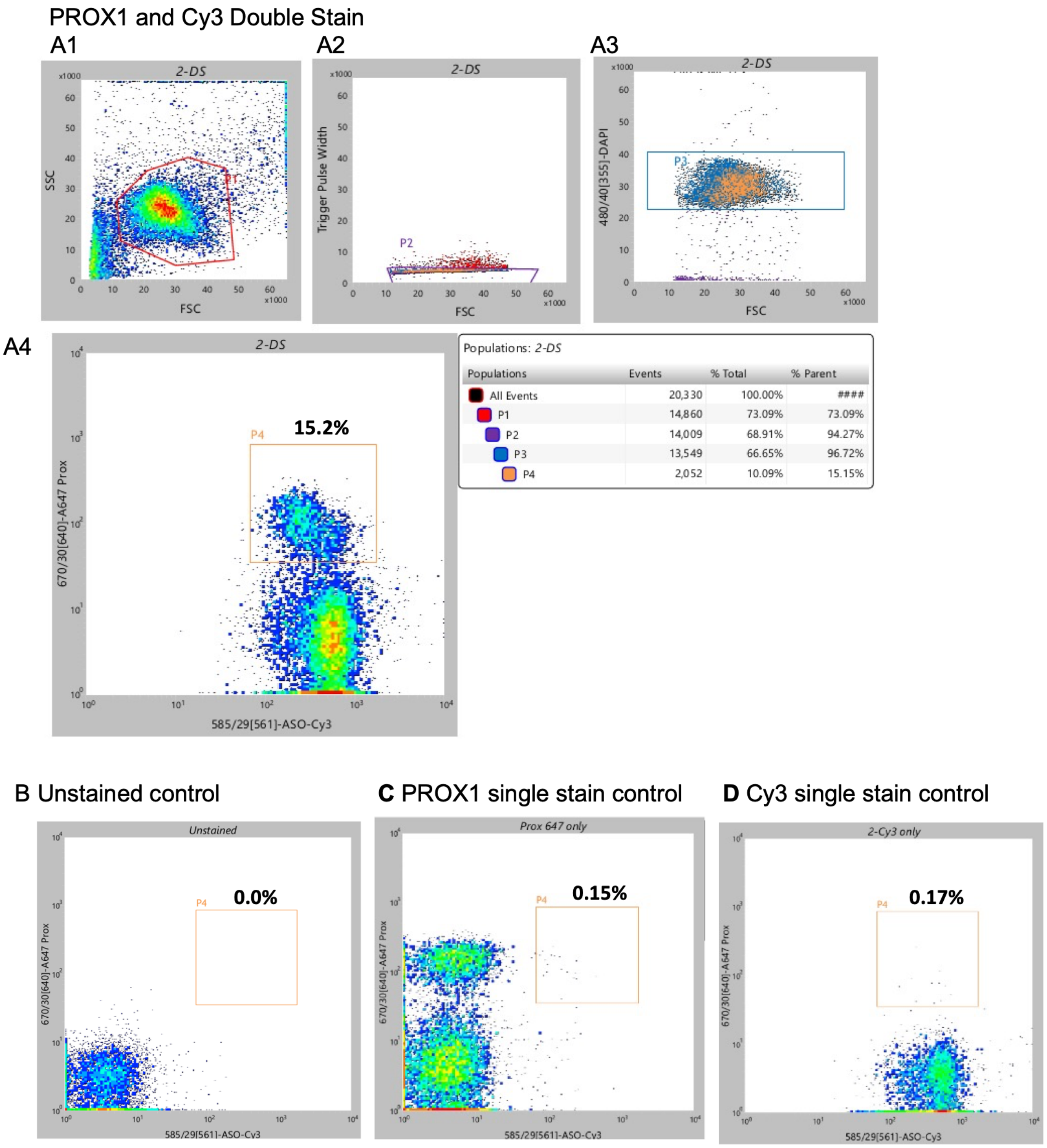


**Extended Data 4.**
**Flow cytometry of nuclei isolated from dorsal hippocampi of mice injected three days earlier in the DG with Cy3 ASO**. (**A**) FSC vs. SSC plot shows a distinct, well-separated population of nuclei (P1 in A1). Applying trigger channel threshold to exclude background from the population of nuclei (P2 in A2). Filtering for DAPI+ nuclei population (P3 in A3). Separation of the Cy3 PROX1+ population (mostly DGCs) from PROX1- cells (P4 in A4). (**B-D**) Background noise in the PROX1+ Cy3 population (yellow box from P4 in A4) is minimal.

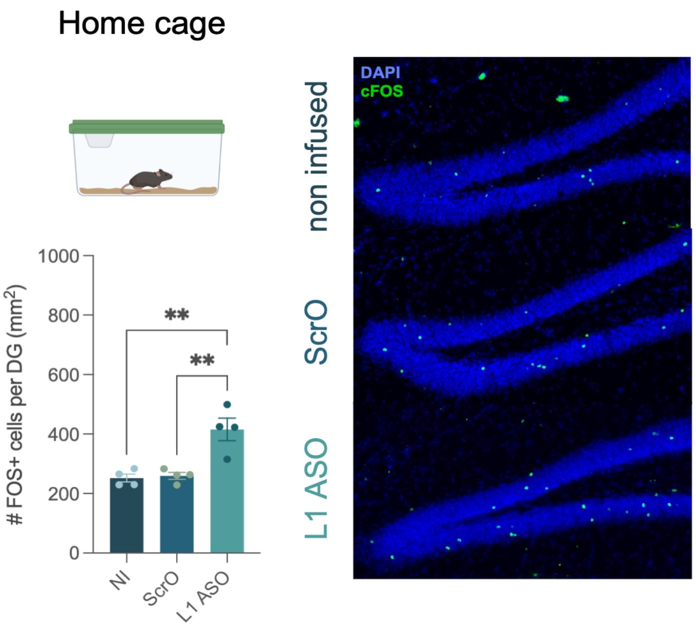


**Extended Data 5.**
**ASO induced increase in the number of FOS⁺ DGCs activated in the home cage 7 days after oligonucleotide delivery**. One-way ANOVA, $F(2,9) = 14.69$, $P<0.0015$. Tukey, $^{**}P<0.005$, N=4/group.

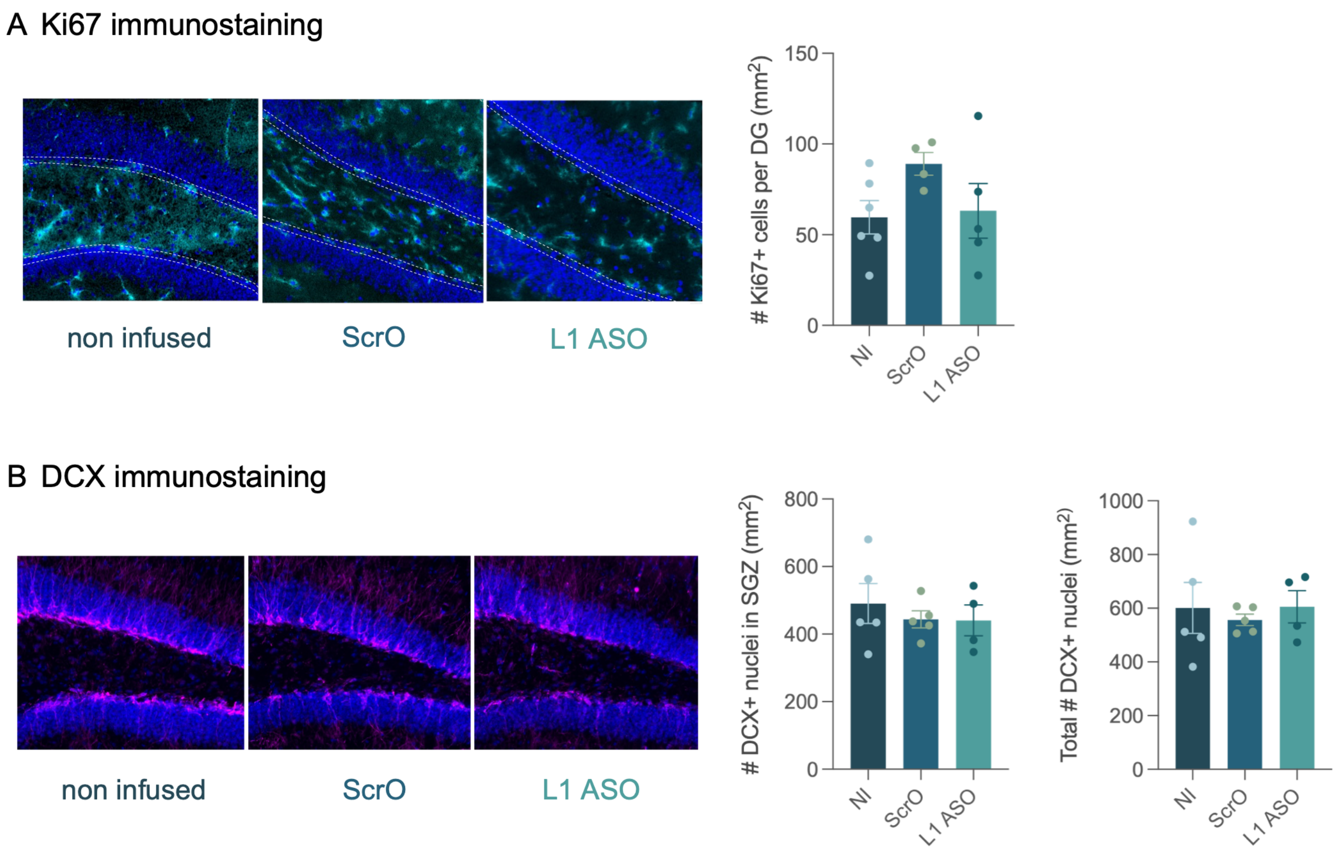


**Extended Data 6.**
**ASO had no effect on the number of ki67+ (A) and DCX positive (B) cells 3 and 7 days post-injection, respectively**. (**A**) Left: Representative images, dotted lines show the boundaries of the SGZ. Right: Quantification of Ki67+ cells in the SGZ. One-way ANOVA, F (2,12) = 1.88, P=0.200, N=6 NI, N=4 ScrO, N=5 ASO. (**B**) Left: Representative images. Right: Quantification of SGZ and total DCX+ cells. SGZ: one-way ANOVA, F (2,11) = 0.39, P=0.688; Total: one-way ANOVA, F (2,11) = 0.17, P=0.849; N=5 NI, N=5 ScrO, N=4 ASO.

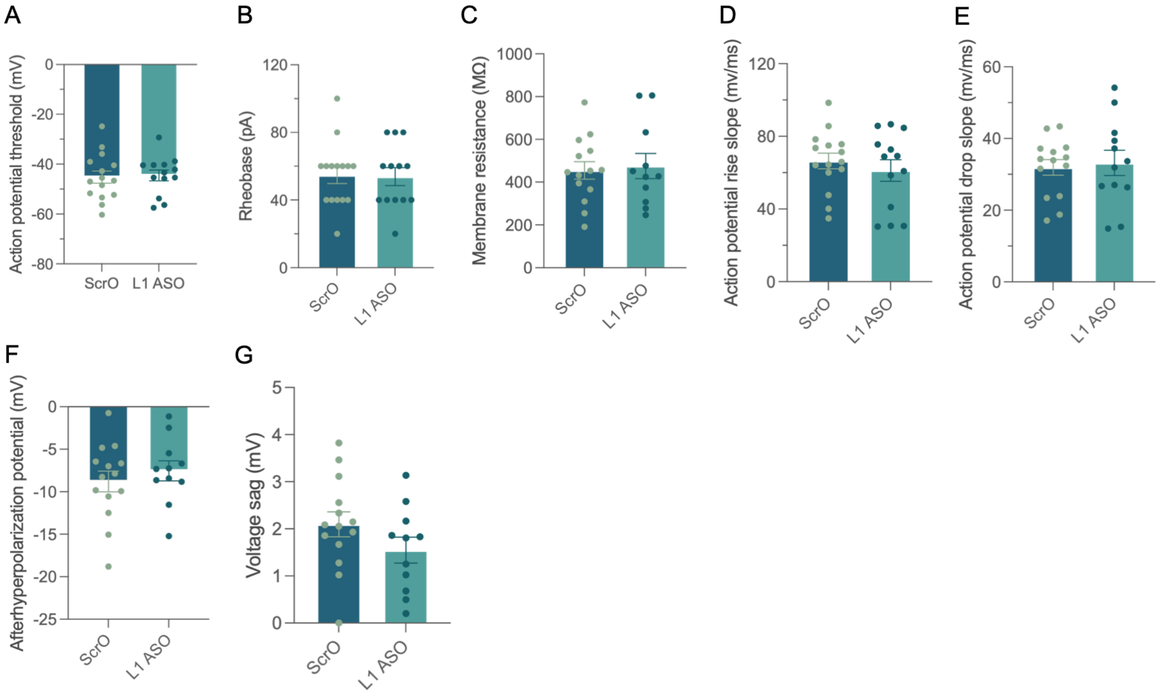


**Extended Data 7.**
**Additional electrophysiological properties of ASO and ScrO DGCs**. **A, B.** ASO and ScrO DGCs had similar AP thresholds (A; unpaired two-tailed t-test: $t_{26}$=0.84, P=0.507). and rheobases (B; unpaired two-tailed t-test: $t_{26}$=0.91, P=0.911). **C.** The membrane resistance between ASO and ScrO DGCs was not different (unpaired two-tailed t-test: $t_{23}$=0.77, P=0.416). (**D-F**) The kinetics of the first evoked action potential are similar between ScrO and ASO DGCs, including the rise slope (D; unpaired two-tailed t-test: t26=0.47, P=0.369), drop slope (E; unpaired two-tailed t-test: t24=0.75, P=0.177), and afterhyperpolarization potential (F; unpaired two-tailed t-test: t23=0.47, P=0.615). (**G**) Voltage sag measured from a -200 pA hyperpolarizing current step, a measure of HCN channel conductance (h-current), did not differ between ASO and ScrO DGCs (unpaired two-tailed t-test: t23=0.17, P=0.807).

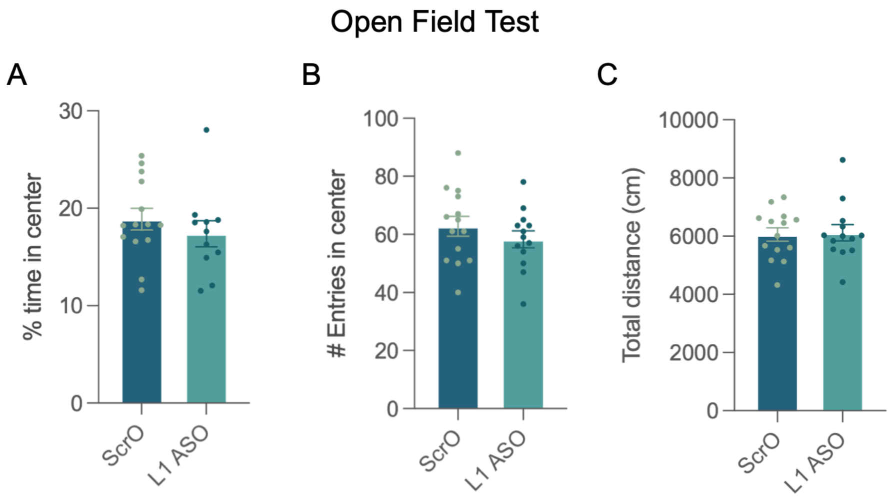


**Extended Data 8.**
**ASO had no effect on anxiety-like behavior in the open field test**. (**A**) Percent time in center. Unpaired two tailed t test, t=0.87, df=23, P=0.391. N=11 (ASO), 14 (ScrO). (**B**) Number of entries to the center. Unpaired two tailed t test, t=0.99, df=25, P=0.331. (**C**) Total distance traveled. Unpaired two tailed t test, t=0.17, df=25, P=0.865.
